# On the innovation and evolution of predatory tactics

**DOI:** 10.1101/530238

**Authors:** Chaitanya S. Gokhale, Anne E. Wignall

## Abstract

Predator-prey systems are ubiquitous across ecological systems. Typical ecological models focus on the dynamics of predator-prey populations. Eco-evolutionary models integrate arms race or Red-Queen like dynamics. The roles of the predator and prey species are always assumed to be static. Nevertheless, sometimes predators can bite off more than they can chew. For example, predators that encounter multiple or dangerous prey types may need to develop new predatory tactics to capture prey. We explore the dynamics of predator-prey dynamics when the prey can injure or kill the predator. This common ecological scenario places pressure on the predator to develop novel predatory tactics to both capture prey and avoid counter-attack from prey. Taking a bottom-up approach, we develop the Holling function mechanistically and then implement it in a model of innovation-selection dynamics inspired by economic theory. We show how an interdisciplinary approach can be used to explain the emergence of complex predatory behaviours. Notably, our study shows why predators may hunt dangerous prey even when safe prey are available. In a broader context, we demonstrate how a multidisciplinary approach combining ecology, evolution and economics improves our understanding of a complex behavioural trait.

## Introduction

Predation is a key selection pressure driving community composition and ecosystem function [1, 2, 3]. Research has previously focussed on the responses of prey in predator-prey interactions [4, 5, 6]. However, there is burgeoning interest in how the behaviour of predators influences population dynamics [7, 8]. There is ample evidence that both within and between-individual variation in predator behaviour influences predator-prey dynamics (e.g. [9, 10, 5, 6]. Our understanding of the degree and forms of influence is still in its infancy despite long recognition of the importance of individual variation [11, 10, 7, 12]. One major source of variation in predator behaviour and hence predation success and population dynamics is individual foraging strategy.

Many predators adopt flexible foraging strategies that incorporate tactics such as stalking, ambush, pursuit or luring to maximise their prey capture success (e.g., [13, 14]). Multiple factors can induce a predator to change foraging tactics, including prey type, environmental factors or satiation levels. For example, in order to ameliorate the risks posed by dangerous prey, predators often adopt specific foraging tactics that are designed to neutralise prey defences. Jumping spiders that prey on other jumping spiders adopt specific cryptic stalking postures that reduce the risk of the prey spider recognising the predator, while adopting non-cryptic stalking behaviour against other prey that is less likely to counter-attack [15]. Similarly, whiptail lizards that prey on scorpions vigorously shake their prey for prolonged periods before consuming them, but adopt a shorter, less violent tactic against cricket prey [16]. The prevalence of flexible foraging strategies in predators are likely to strongly co-vary with the population dynamics of predators and prey.

Predators and prey have some of the closest eco-evolutionary relationships. There is often intense selection pressure on both predator and prey populations to develop and establish novel predatory or anti-predator traits (e.g. [17]). Changes in predatory behaviour can result in significant shifts in predator-prey population dynamics [18, 19]. How predators develop new predatory tactics is not well understood. Innovation in animals (and across scales of organisation) has been a subject of intense investigation in recent years [20, 21]. Here, we develop a model that demonstrates that innovation dynamics can result in the development and maintenance of complex predatory tactics. We then quantify the population dynamics between predators with flexible foraging strategies (i.e., multiple predatory tactics) and their prey.

We develop a framework which is a combination of traditional ecological theory, evolutionary game dynamics and innovation theory [22, 23]. We develop two prey types, safe and dangerous. Predators could have simple strategies employing straightforward hunting tactics, or complex having novel and complicated hunting tactics. We derive the Holling responses for the hunting strategies and the different prey types by using a decision tree based approach inspired by game theoretic analysis. While the population dynamics of such a model can be extensively studied, our aim is to hypothize a mechanism which brings about innovative complex predatory tactics in the first place.

## Model and Results

### 2.1 Population dynamics

Before addressing dangerous prey and novel hunting tactics, let us begin with a simple predator-prey model. The abundance of the prey is given by *x* and that of the predator by *y*. The following set of differential equations can then describe the dynamics,

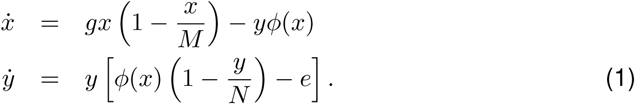

The prey increases in density at a certain fixed rate *g* but the density is capped by the carrying capacity *M*. The prey density is reduced by consumption by the predator according to the function *ϕ*(*x*), a function of the prey density. As such, predators can increase in density while dying at a constant rate *e*. In the simplest case one can assume that *ϕ*(*x*) = *x*. The analysis of such a simple predator-prey system can be found in the classical texts of theoretical biology [24].

Next, we extend the system to include two prey types (safe and dangerous). The two types of prey reflect natural communities in which predators often have multiple prey types, rather than specialising on just one. We track the abundances of safe and dangerous prey by *x*_*s*_ and *x*_*d*_ respectively. Thus now the system would be made up of the two prey types and a simple predator (See SI.1 for the dynamical equations).

Let us imagine a predator which can assess the risk of attacking a dangerous prey and adjust its behaviour accordingly. We assume that both prey (predator) types fall in the same niche such that they compete for space and therefore have a combined carrying capacity of *M* (*N)* respectively. Thus we can track the population dynamics of the prey *x*_*s*_ (safe), *x*_*d*_ (dangerous) and the predator *y* (simple) and *z* (complex) (see Table in SI.1). Both the simple and complex predatory strategies, as well as the safe and dangerous prey types, can co-exist. However, how does the complex strategy arise? To understand the origin of such strategies we first start by quantifying the decisions made by predators during their interactions with prey.

### 2.2 How do predators make decisions?

To understand how new predatory behaviours can come about, first, we need to understand how the predators progress through a hunt. The response function usually assumed to be merely a linear function of the prey density may not be linear. How does the density of prey influence prey capture rates? Holling [25] addressed this question resulting in the now-famous Holling response curves. The response curves from Holling’s experiment and their variations have been derived and readdressed to take into account various eco-evolutionary factors such as multiple prey types, different types of prey or including prey defence mechanisms [26, 27, 28]. While we could adopt one of the Holling responses developed, we would lose any mechanistic interpretation of the terms involved. We derive the response function using a decision tree technique from game theory [22]. This decision tree model helps us step through the decision-making processes of predators.

#### Simple predator

A simple predator searches for prey, spending time *t*_*s*_ and encounters the two prey types, safe and dangerous, with probabilities proportional to their relative abundances in the population. The probability of encountering safe prey is thus *S* = *x*_*s*_*/M* and of encountering dangerous prey is *D* = *x*_*d*_*/M*. Empty patches are thus encountered with probability 1 *-S - D*. A predator that encounters safe prey follows the left side of the decision tree whereas for dangerous prey it follows the right side (Figure. 1). The payoff of each set of decisions is provided at the bottom of the leaves. The time required to reach that decision is on the left for safe prey and on the right for dangerous prey encounters. Since the prey handling time depends on prey type, these times can differ.

**Figure 1:**
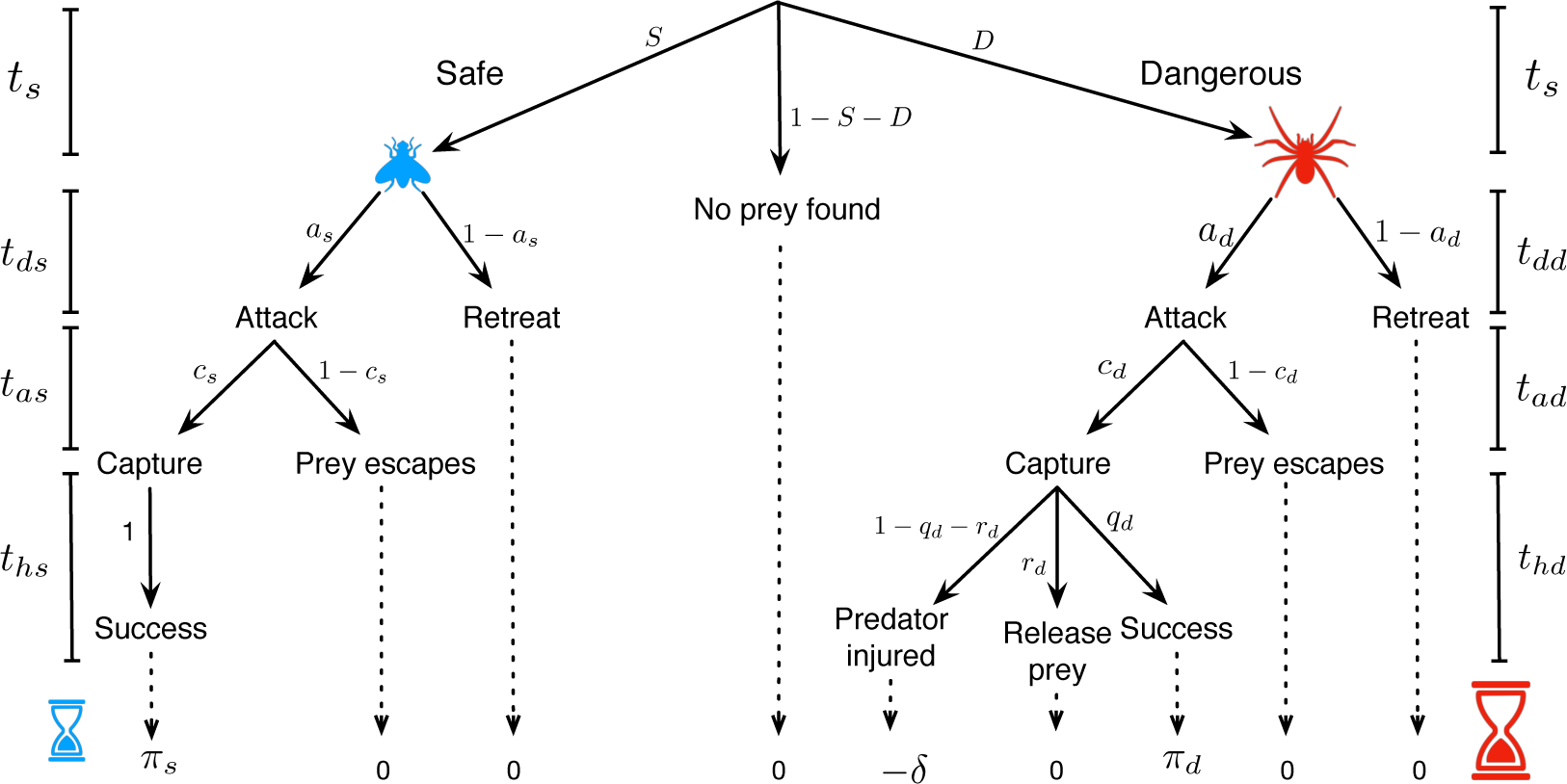
Decision tree for a simple predator. The predator encounters safe prey with probability *S* = *x*_*s*_*/M* and a dangerous prey with probability *D* = *x*_*d*_*/M*. The probability that there is no prey in the patch follows 1 - *S - D*. The predator first decides to attack with probability *a*_*s*_ (*a*_*d*_). If the predator chooses to attack, the capture probability is denoted by *c*_*s*_ (*c*_*d*_). However, after prey capture the predator can decide whether to release the prey (finding it to be too dangerous) with probability *r*_*d*_ or it may be injured in the process of attacking with probability 1 - *q*_*d*_ - *r*_*d*_ where *q*_*d*_ is then the eventual probability of successful capture of dangerous prey. For safe prey, these probabilities of predator injury do not appear. Each of these steps requires a certain amount of time, given on the edges of the tree with the notation *t*. The outcomes of following each branch of the decision tree are collated at the terminus of each branch.

As shown in Figure 1, there are nine possible outcomes (*O*_1_ to *O*_9_). Each outcome has a probability given by the product of the probabilities leading to that outcome. For example, the probability of encountering a safe prey, then deciding to attack it, but then the prey escaping is given by *x*_*s*_*O*_2_ where *O*_2_ = *a*_*s*_(1 - *c*_*s*_).

The functional responses of the simple predator upon encountering the two prey types are then denoted by,

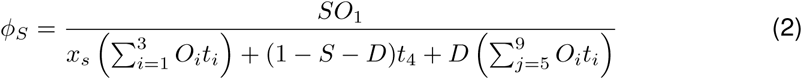

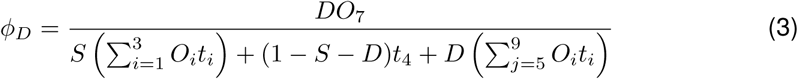

which simplify to,

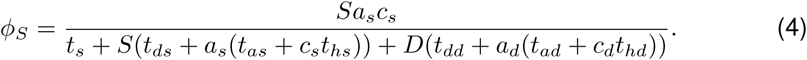

and,

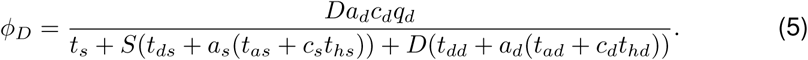

These functional responses look unfamiliar because we have assigned both prey types to be present. As in [22], we can recover the traditional form of the Holling response by normalising the search time *Mt*_*s*_ = 1 and focusing on cases when only one prey type is present (either safe (*x*_*s*_ = *M)* or dangerous (*x*_*d*_ = *M)*). These are the assumptions which the traditional Holling response is based on, and our functions become,

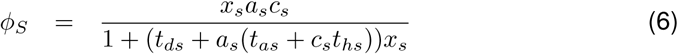

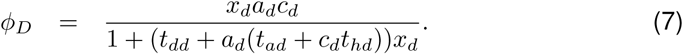

We assume that the simple predator cannot differentiate between the safe and dangerous prey. Therefore it decides to attack any prey with the same probability (*a*_*s*_ = *a*_*d*_ = 0.9) and attacks instantaneously (*t*_*s*_ = *t*_*d*_ = 0). However the dangerous prey is harder to capture (*c*_*d*_ = 0.2 *< c*_*s*_ = 0.8). The difference in the timing lies in the handling time. For different magnitudes of handling times for the dangerous prey, we see that the capture rate of safe prey is always higher than that of dangerous prey Fig. 2 (left panel).

**Figure 2:**
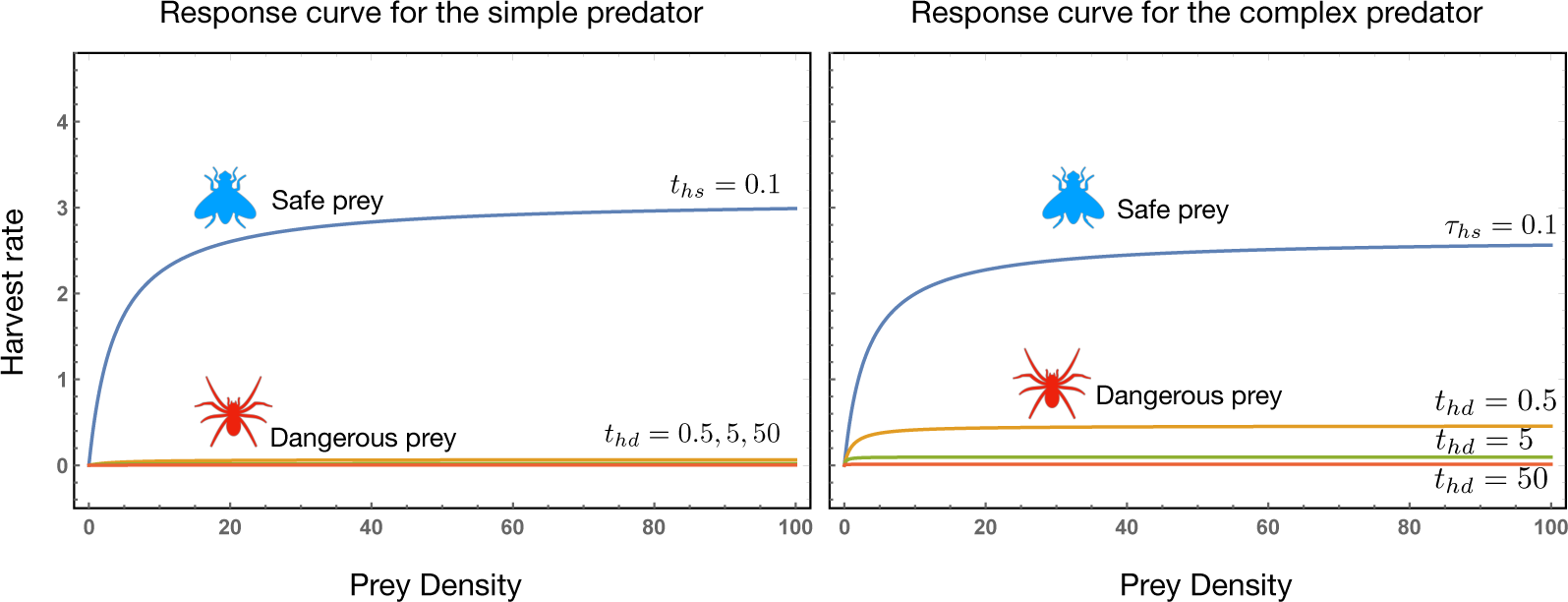
Left panel. For the simple predator the chance of hunting the safe prey depend on *as* = 0.9 and *cs* = 0.8. Similarly, successfully hunting a dangerous prey depends on *a*_*d*_ = 0.9, *c*_*d*_ = 0.2, i.e. the predator does not differentiate between the prey types when deciding to attack *t*_*ds*_ = *t*_*dd*_ = 0.0 but the capture probabilities differ. Furthermore *q*_*d*_ = 0.1 and *r*_*d*_ = 0.5, i.e. there is only a small chance of success as there is a higher risk of the predator getting injured. The handling time for dangerous prey *t*_*hd*_ = 0.5, 5, 50 is more than handling safe prey *t*_*hs*_ = 0.1 resulting in a progressively lower prey capture rate. **Right panel.** For a complex predator, the decision tree is similar however we have *a*_*s*_ = 0.9 whereas *a*_*d*_ = 0.8, i.e. the predator discerns between the two prey types and can actively choose to attack the dangerous prey less often than the safe prey. A complex, novel tactic can result in a higher capture success with *c*_*d*_ = 0.9. However an innovative strategy requires an investment of time, with *t*_*dd*_ = 0.1. The invested time and complexity result in a higher chance of success *q*_*d*_ = 0.55*, r*_*d*_ = 0.1 with a smaller chance of injury. For both scenarios we set *ts* = *τs* = 1. The resulting prey capture rates follow a similar pattern to that predicted in [29].

Thus using a decision tree model coming from game theory, we have provided a mechanistic basis for the functional response of a simple predator to dangerous prey. It is possible to analyse the response dynamics further using the static concepts of equilibrium selection as is done extensively in [22] for safe prey. However, our aim here is to analyse the functional response of a predator employing complex strategies, and subsequently, the dynamics of predator-prey interactions.

#### Complex predator

A complex predator assesses the risk posed by safe and dangerous prey (Fig. 3). First, the complex predator needs to identify the kind of prey it has encountered. Risk assessment results in a delay in making an initial decision to attack or abort the hunt as compared to the simple predator *τ*_*dd/ds*_ *> t*_*dd*_. Next, the predator selects a direct attack tactic for safe prey, while for dangerous prey a different tactic is employed.

**Figure 3:**
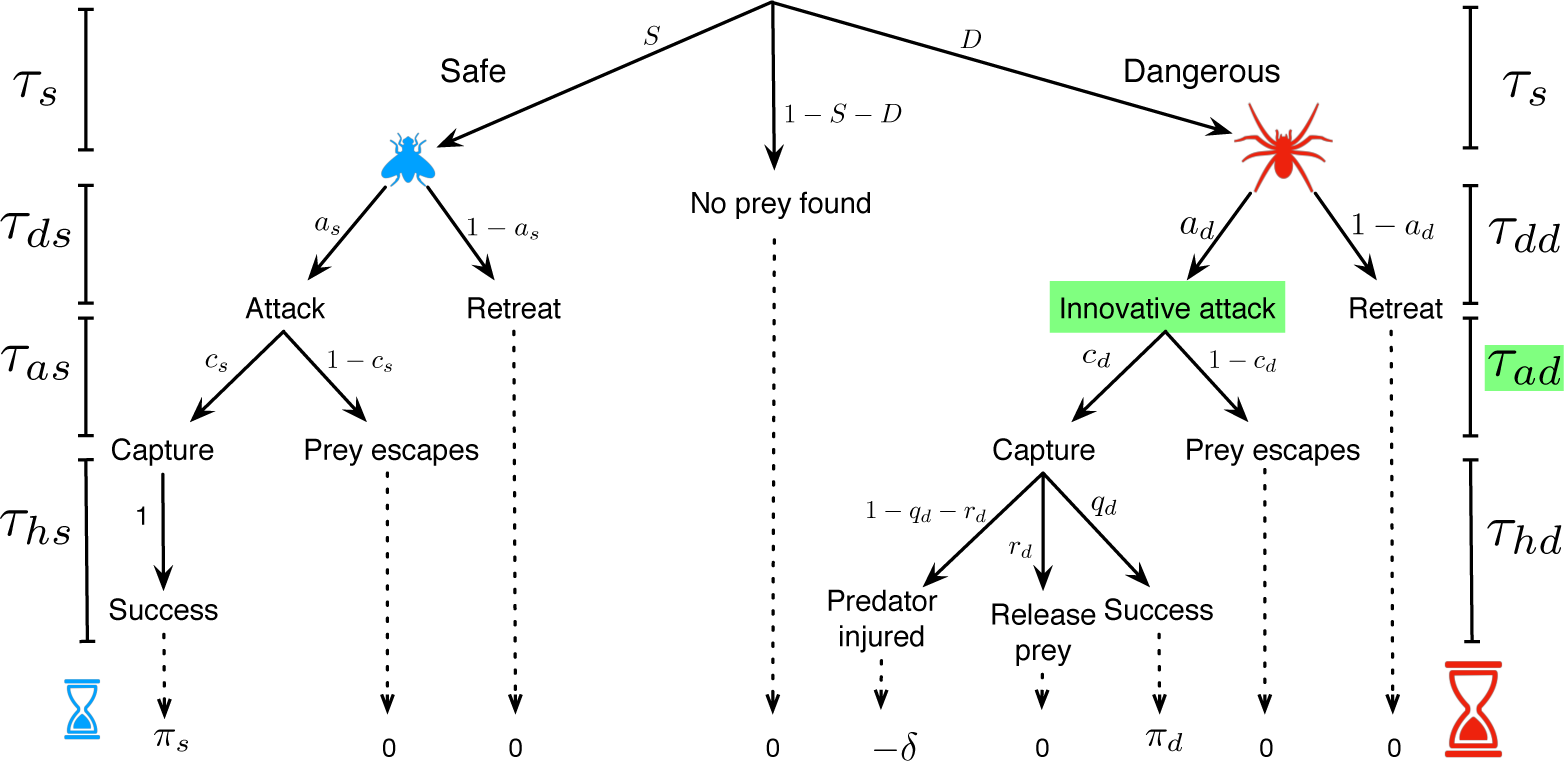
Decision tree for a complex predator. The predator encounters safe prey with probability *S* = *x*_*s*_*/M* and a dangerous prey with probability *D* = *x*_*d*_*/M*. The probability that there is no prey in the patch follows 1 - *S - D*. For the safe prey the dynamics proceeds in a similar manner as that for the simple predator (Fig. 1). When deciding to attack the prey, the complex predator can distinguish if the prey is safe or dangerous. This process of discerning takes time such that as compared to the simple predator we have *τ*_*dd/ds*_ *> t*_*dd*_. Given a dangerous prey the predator chooses to employ a novel innovative attack technique. The execution time of the innovative attack will depend on the exact type of the innovation. The structure of the rest of the decision tree is again the same as in Fig. 1.

Using the same analysis technique as for the simple predator, we can write down the response curves for the complex predator when feeding on the two prey types as,

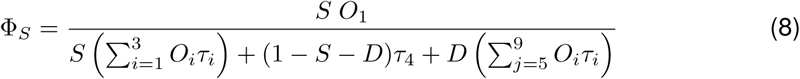

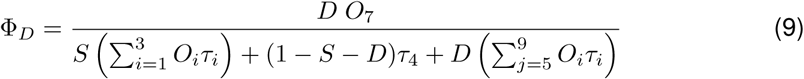

Since the complex predator has invested time in assessing the difference between the prey types, it attacks the two prey types with different probabilities (say *a*_*s*_ = 0.9 *> a*_*d*_ = 0.8). Employing a novel hunting technique specific to the dangerous prey results in a higher capture probability as compared to that of a simple predator. Subsuming the costs in the amount of time spent in the discriminatory behaviour, the complex predator overall performs worse at capturing prey although it can capture the dangerous prey better than the simple predator (compare panels in Fig. 2).

While we have characterised the response curves of the predators to the different prey types, they are not in the same currency as that of energizable metabolic units. The energy gain for the predators (simple and complex, *f* and *F)* is denoted by,

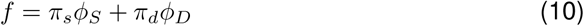

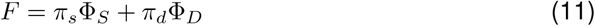

When a predator attacks dangerous prey, the rate at which it can get injured is a function of the density of the dangerous prey, just as a Holling response. We capture injury rates with *ϕ*_*I*_ and Φ_*I*_ for the simple and complex predators respectively calculated in a manner similar to Eqs. (7),(9).

### 2.3 Population dynamics revisited

Now that we have a mechanistic description of the functional responses we can revisit the population dynamics of the two prey types, safe and dangerous, and the two predator types, simple and complex. The dynamics follow the following set of equations,

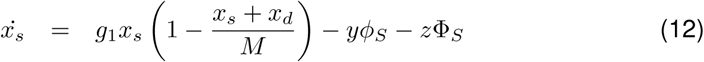

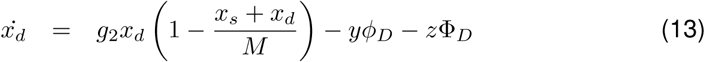

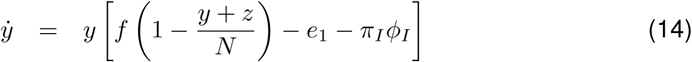

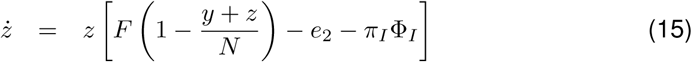

The growth of the two prey follows a logistic pattern whereas their deaths feed the predators. The predators die with an intrinsic death rate but also after suffering injuries from encountering the dangerous predator. The population dynamics of such a system is akin to the dynamics of two prey and two predators [30, 31].

So far, we have assumed that the complex predator is present without discussing its origins. At the core of this study lies the question of how a complex predatory strategy can come about in the first place. We thus discuss the different ways a predator can “innovate” a complex novel predatory tactic.

### 2.4 How do predators innovate?

Until now we have followed the population dynamics assuming that ‘somehow’ an innovative predatory tactic comes about and then spreads in the population. However, how exactly does this innovative tactic arise? We assume that a predator can employ different types of simple strategies a set we define as the core tactics. A complex predator can also have a set of innovative predatory tactics a set we define as the ‘novel’ tactics. Here, we define a route between the set of simple core tactics and the complex novel tactics. To do so, we make use of the concept of innovation.

In engineering and economics, the concept of innovation has been studied extensively [32]. Economics defines three conditions that need to be satisfied to drive innovation [33] – a recognised need, competent people with the required technology and, financial support. We argue that similar factors drive innovation in biology,

- a positive selection pressure,
- prerequisite genetic, phenotypic, behavioural variability,
- within bioenergetic and physicochemical bounds.

#### Core tactics

We assume that there are a few core tactics that a predator can utilise. Core hunting tactics are the basic techniques which a predator possesses to hunt “safe” prey, i.e. prey which does not pose any risk of injury or death to the predator itself. The predator can switch between these tactics with a switching rate *µ*. For our model, we characterise three core tactics. Ambush (A), Lure (L) or Pursuit (P) could be such core tactics as visualised on the left in Fig. 4.

**Figure 4:**
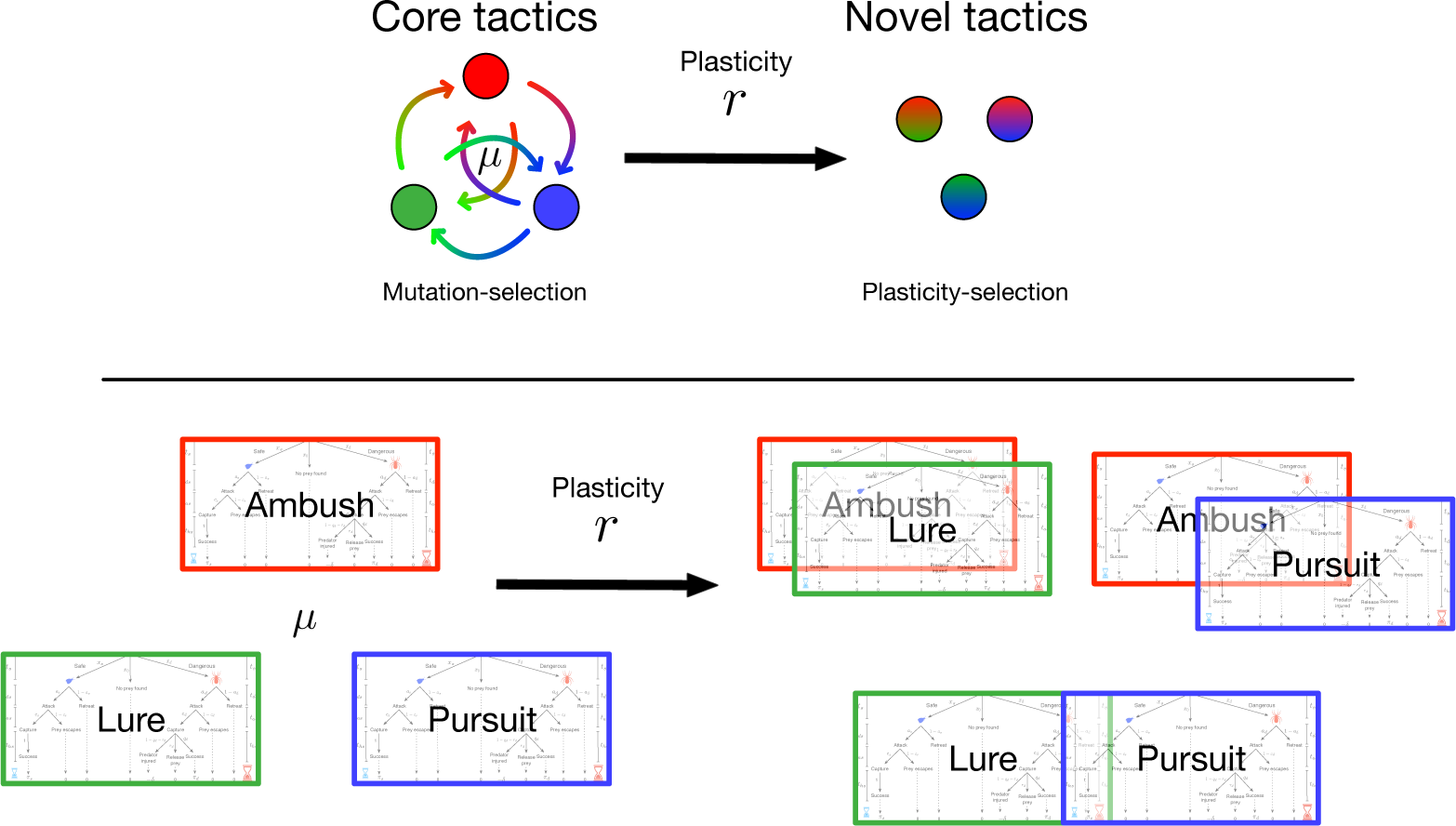
Switching-plasticity dynamics. Evolving complex novel strategies is assumed to be possible from a given set of simple, core tactics. If a predator has the tactics of ambush, lure and pursuit in its basic repertoire then it is assumed that it can switch between the three with a probability *µ*. With probability *r* it is possible to recombine two of these ‘simple’ core tactics into a blend. The new blended tactic is termed the ‘complex’ novel tactic. Thus, it is then possible for a predator to Lure and Ambush, Ambush and Pursuit or Lure and Pursuit.

#### Core to novel

To see how novel strategies can arise, we take inspiration from evolutionary game theory. The core tactics could switch between themselves, but since they are behaviours, they could also combine. For example, a predator may wait to ambush prey with a lure out, a typical fishing strategy (e.g. [34]). This ability comes with a well-justified tradeoff [35] as the behaviour requires either a large brain or an investment in physical structures or substances to attract prey. All these costs could be captured by an overall higher death rate of these innovative tactics.

Thus the overall cost of this innovation reduces the prey capture rate of the predators employing the novel tactics *z*. In a broad sense, the predators thus have two strategies, simple or complex. Simple strategies involve using core tactics while a complex strategy involves the use of the novel, innovative tactics. In general for *n* core tactics we have 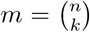 possible “innovative” hunting tactics. The total number of tactics in the spider population are thus *n* (core tactics) + *m* (derived novel tactics). The dynamics are now further modified as,

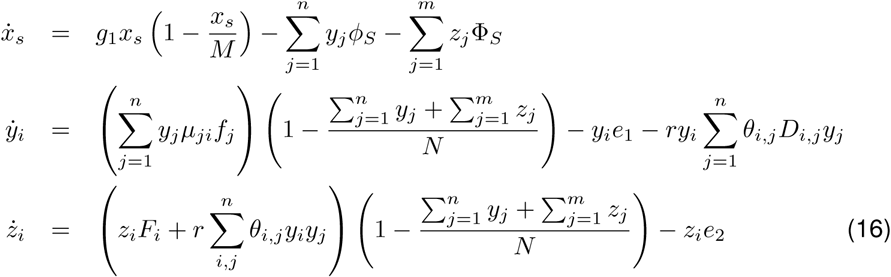

where all the core tactics are denoted by *y* and the novel tactics by *z*. Two core tactics blend with a rate *r* and the elements of the matrix *D* determine the contribution of the individual core tactics in forming the new tactic. We have *D*_*k,k*_ = 0 and *D*_*i,j*_ + *D*_*j,i*_ = 1. The binary operator *θ*_*i,j*_ which can be either 1 or 0 dictates whether a certain combination is even possible or not, for e.g. *θ*_*i,i*_ = 0. Even if there is a continuous influx due to the recombination rate *r*, the novel hunting tactics have a reduced prey capture rate of the safe prey as compared to the core tactics. This cost does not allow the complex tactics to flourish even if they originate.

### 2.5 Full model

To build up the model stepwise we now include the dangerous prey as well. Denoted by *x*_*d*_, as before, the initial abundance of this prey type is assumed to be extremely low. This property can be captured by the now complete model as given by the equations:

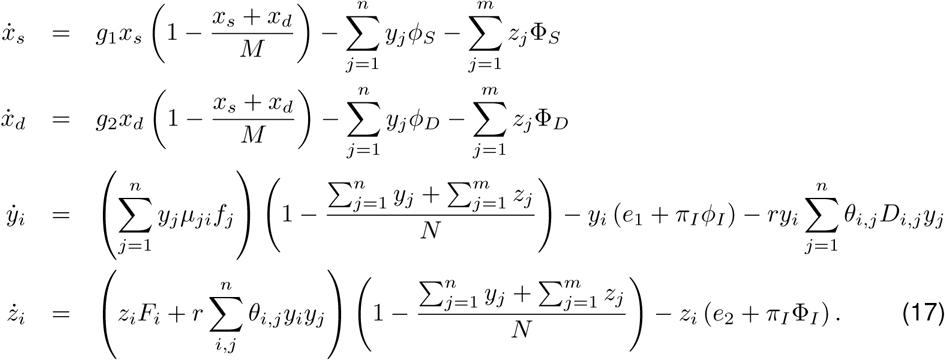

While it is straightforward to write down the dynamics of all the types in the population and possibly further analyse the model, we focus on the definitions of the terms within this theory. For example, we have calculated the response functions for a simple and a complex tactic in Fig. 2. Now, we have three simple core strategies, and the example harvest rates for these denoted in Fig. 5 (left panel). A binary combination of the core tactics results in the three complex novel tactics. The example response functions are illustrated in Fig. 5 (right panel) for each complex novel tactic.

**Figure 5:**
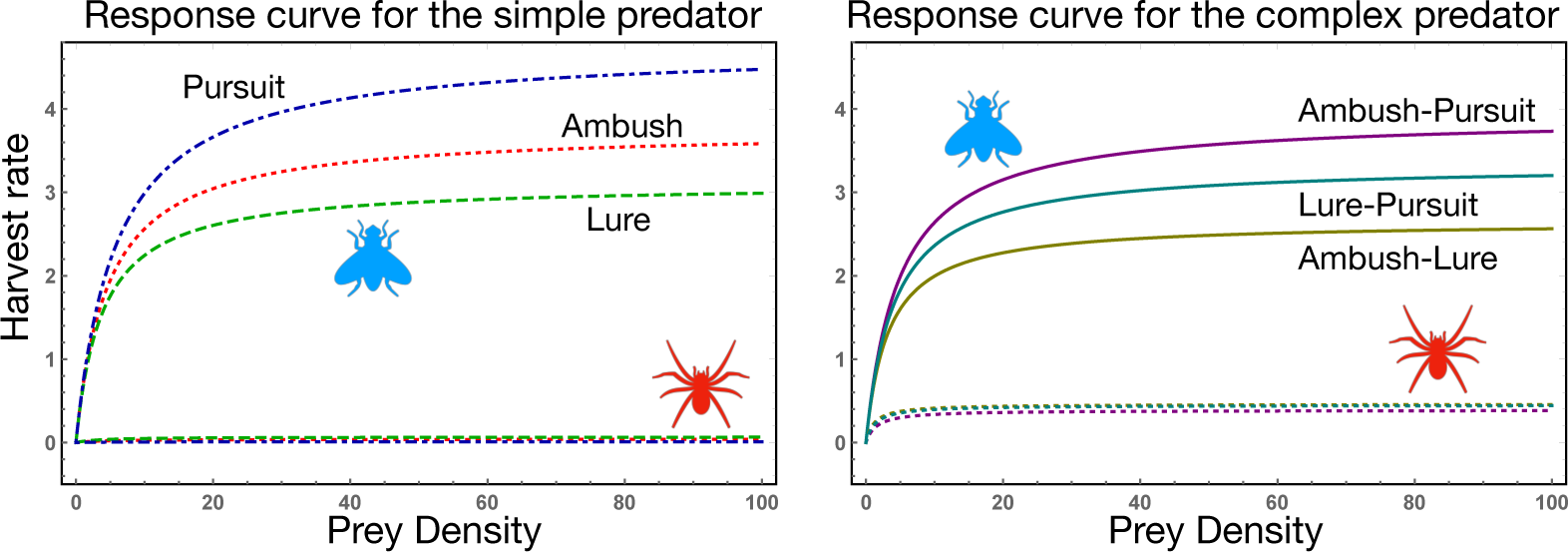
Harvest rates for the core and novel strategies. For the simple predator and the complex predator some of the parameters have the same values. For example we have *t*_*s*_ = *τ*_*s*_ = 1, *a*_*s*_ = 0.9, *c*_*s*_ = 0.9 *t*_*ds*_ = *τ*_*ds*_ = 0, *t*_*hs*_ = *τ*_*hs*_ = 0.1 and *t*_*hd*_ = *τ*_*hd*_ = 0.5. The difference lies in the time taken to decide whether to attack a dangerous prey or to retreat *t*_*dd*_ = 0 since the prey needs to be identified as dangerous. The individual attack tactics also have their own parameter settings, for the simple predator the tactics are Ambush (*t*_*as*_ = 0.15, *t*_*ad*_ = 0.15, *q*_*d*_ = 0.1 and *r*_*d*_ = 0.8), Lure (*t*_*as*_ = 0.2, *t*_*ad*_ = 0.2, *q*_*d*_ = 0.1 and *r*_*d*_ = 0.6) and Pursuit (*t*_*as*_ = 0.1, *t*_*ad*_ = 0.1, *q*_*d*_ = 0.05 and *r*_*d*_ = 0.9). For the complex tactics we have, Ambush-Lure (*a*_*al*_ = 0.8, *c*_*al*_ = 0.9, *τ*_*as*_ = 0.25, *q*_*al*_ = 0.55 and *r*_*al*_ = 0.1), Ambush-Pursuit (*a*_*ap*_ = 0.8, *c*_*ap*_ = 0.8, *τ*_*as*_ = 0.14, *q*_*ap*_ = 0.4 and *r*_*ap*_ = 0.6) and Lure-Pursuit (*a*_*lp*_ = 0.8, *c*_*lp*_ = 0.85, *τ*_*as*_ = 0.18, *q*_*lp*_ = 0.5 and *r*_*lp*_ = 0.15). The space of success of these strategies is discussed further in Fig. 6. The time required to execute the the complex tactics *τ*_*ad*_ are assumed to be A-L (*τ*_*al*_ = 0.5), A-P (*τ*_*ap*_ = 0.3) and L-P (*τ*_*lp*_ = 0.4).

How tactics combine depends on the organism and the type of tactics. If we imagine two straightforward hunting behaviours then perhaps what combines is not the actual predatory tactic, but the phenotypic adaptations developed for each tactic. For example, the ability to use wind direction to avoid detection by prey plus to be able to map the three-dimensional surroundings to approach prey cannot be combined if the appropriate phenotypic adaptation to detect wind movement and three-dimensional optic machinery have not evolved in the same animal. While our concept of combining tactics is a simplification over the natural processes of functional overloading, it allows us to make inroads into theoretically unchartered territory. For example, we can map the complex novel tactics of a predator together with the simple core tactics when attacking a dangerous prey. The tactics can be visualised by measuring the mutually exclusive probabilities of hunting success, predator injury and (dangerous) prey release in a probability space as in Fig. 6. Core tactics are predicted to be the least successful predatory tactics, but require the least amount of investment by the predator. Novel tactics are expected to reduce the risk of predator injury while increasing prey capture success. However, the cost of novel tactics (e.g., via time investment) is higher than core tactics (Fig. 5). In our example, tactics that include luring prey, which requires behavioural modification of the prey (i.e., the prey must approach the predator) are the most risky for the predator and yet are the most likely to result in prey capture.

**Figure 6:**
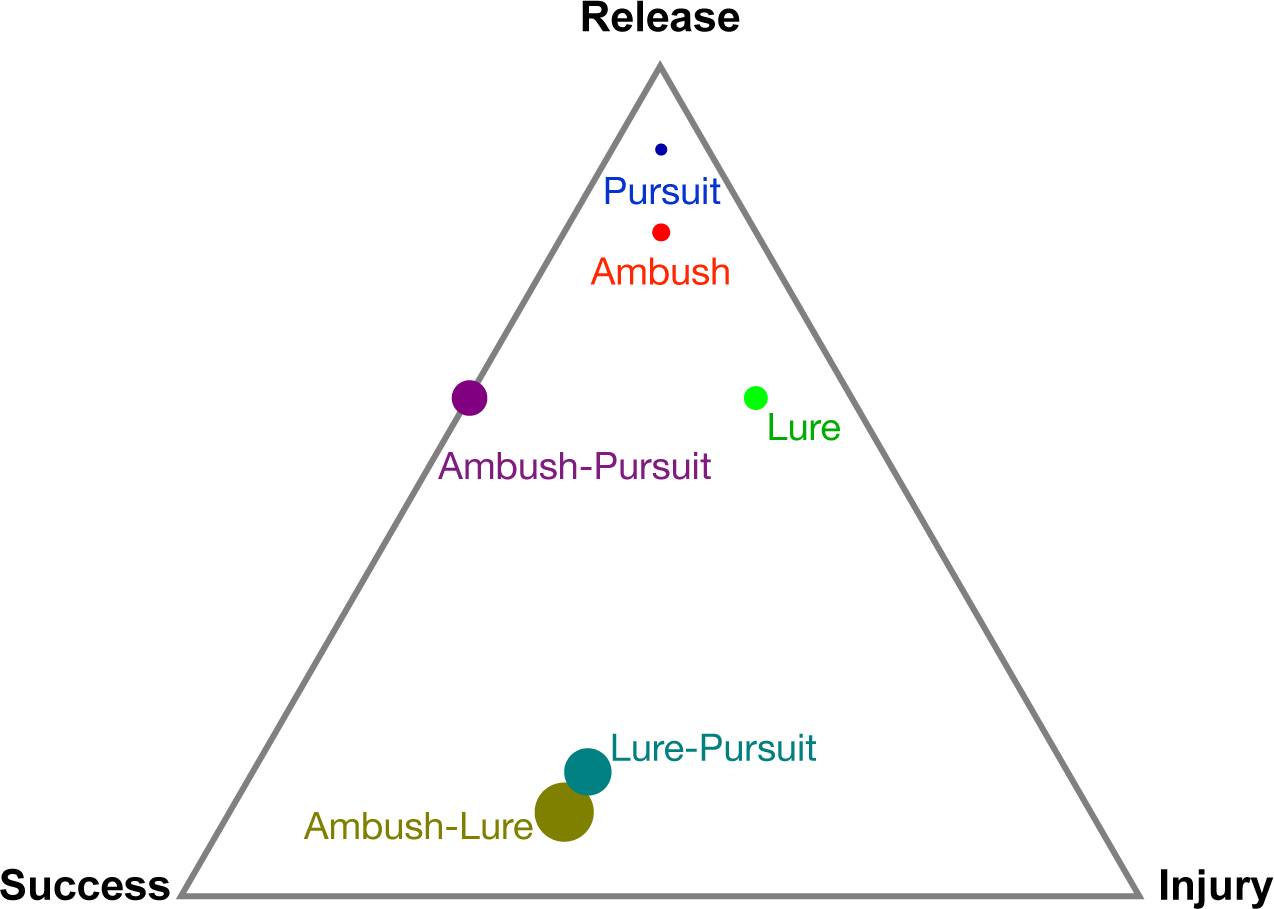
Result space for core and novel predatory tactics. Each predatory tactic is defined by its Success, Injury and Release probabilities. We can thus plot the tactics in this space where the vertices result in the labelled outcome being realised with probability 1 (and with probability 0 at the opposite edge). Besides these three probabilities, each tactic also takes its own time for execution. The size of the points represents this. Ambush-Lure is the most time-consuming tactic while pursuit is the fastest.

We have thus completed the development of the full model. Starting with the classical ecological model of predator-prey dynamics (Fig. 7 (a)), we added dangerous prey in the model (b). This resulted in loss of the stability of the classical predator-prey cycles. Although suffering from a lower growth rate compared to safe prey, dangerous prey can deter the predators and thus leads to a dominance of dangerous prey over other types. Then we included multiple simple ‘core’ tactics and allowed the predator to switch between them (c). While this allows multiple predatory tactics to be exploited, some performing better than others, they are still simple. The dangerous prey can evade such tactics and maintains its superiority. Finally allowing for innovation, complex ‘novel’ hunting tactics arise (d). The novel tactics, whose functionality until now lay dormant, can now invade the hunting strategy space and co-exist together with the ‘core’ tactics. For the illustrated parameter, we observe a re-emergence of predator-prey cycles (at least in the short run) before resulting in a static but stable antagonistic co-existence between the (simple and complex) predators and (safe and dangerous) prey types. While Ambush was the most successful when only safe prey was present, in a mixed situation where both prey types are present, a combination of Luring and Pursuing works best.

**Figure 7:**
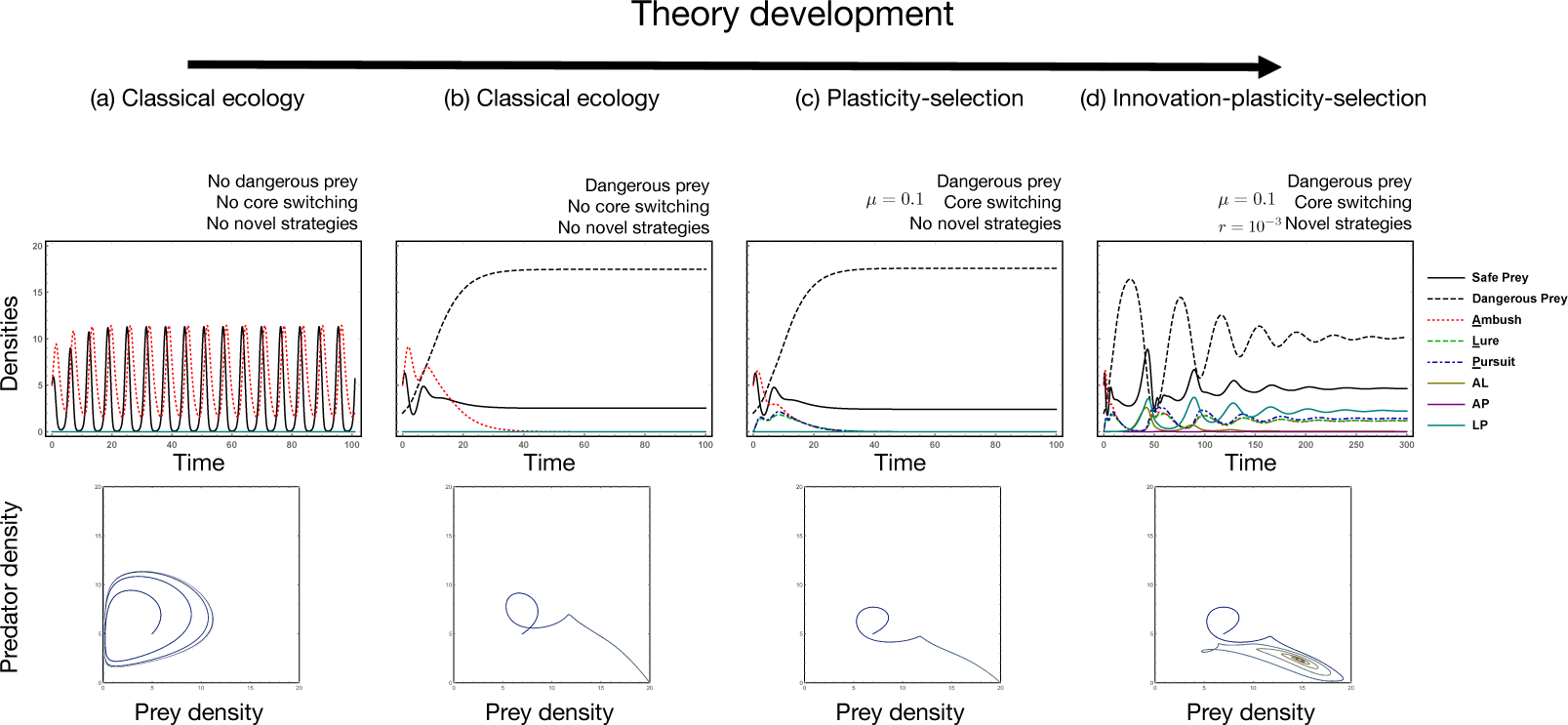
Illustrative example of the full model. The full model as described in the set of equations in Eqs. 17 is illustrated here for four different scenarios. From the left to right we illustrate how we have proceeded with the theory development. (a) A simple one predator-one (safe) prey system is described which provides us with the classical Red-Queen dynamics. (b) The addition of a dangerous prey, (which can harm the predator) can result in the extinction of the predator and a coexistence of the prey types. (c) If we allow for multiple predatory hunting tactics (all core tactics), with a switching rate of *µ* = 0.1 we see that the tactics are explored by the predator however since they all are dominated by the dangerous prey, eventually they all die out. (d) Together with switching (*µ* = 0.1) if complex and novel predatory tactics can evolve (at rate *r* = 10^-3^) we see that the novel tactics can increase in abundance in the population and coexist with different hunting tactics. We have used the same parameters for the tactics as described in Fig. 6 which shows that the A-L tactic has the highest success probability. However, the temporal dynamics reveal that the successful novel tactic, L-P, emerges via an evolutionary method rather than only success maximisation.

## Discussion

Animals, from invertebrates to humans can show innovative behaviour [20]. Innovation can even be applied from cells to complex societies [21]. We present a method of incorporating innovative behaviour in classical predator-prey dynamics by testing how new predatory tactics are developed and maintained in populations. Innovative traits such as those that arise from plasticity could eventually come under developmental control when in the right ecology [36].

A simple model of predator-prey dynamics forms the basis of our model. However, we uniquely derive the prey capture rates of the predator using a decision tree like approach pioneered by [22]. This bottom-up approach allowed us to expand the model to include dangerous prey and complex predators who can employ novel hunting tactics. Probing further, we have addressed how such novel tactics can arise in the first place.

Technological innovation can be studied by making use of the same dynamical systems as used in evolutionary dynamics [37]. An innovative predator can formulate alternative hunting tactics. As it is more parsimonious to tweak already available material, we assume that the novel hunting tactics could be a blend of the already present ‘simple’ core hunting tactics. We thus demonstrate how novel and complex tactics that one can observe in predators can originate from a few core simple tactics. Evidence for this comes from *Portia* jumping spiders that use trial and error to derive vibrations to attract their web-building spider prey within range of attack [38]. Jumping spiders make an excellent model system for examining the processes of behavioural innovation in a predator due to their excellent vision and complex cognitive abilities in a small brain [39, 40]. Further, jumping spiders have multiple predatory tactics including ambushing, luring and pursuing prey [41, 42]. These predatory tactics are often adapted for prey type and context [43, 44].

Of further interest would be the rate at which novel tactics arise in the population. Factors that influence the probability of novel predatory tactics arising are likely to include animal personality and changing communities (either through animal introductions or range expansions) providing new sources of prey. Predator personality has already been demonstrated to affect prey capture rates [45, 46, 47]. The effects of personality are likely to extend to predatory tactics. Once available phenotypically, such tactics could in principle come under developmental control [36]. In our model, we assumed that once such innovative tactics appear in the population, then they can be passed along genetically. The rapid evolution of behavioural tactics in a predatorprey arms race between a spider and fly indicates that this is a reasonable assumption [17]. If the inheritance of predatory tactics were strictly social, then we would need to include hybrid modelling techniques. That is, after every generation, the novel predatory tactics would be removed from the population. However, if generations are over-lapping, we could also envisage a role of predator learning in the transmission of predatory tactics. Furthermore, a difference in the lifespan of the predator and prey could also facilitate within generation learning opportunities the role of experience. Certainly experience influences predatory behaviour (e.g., [48]). Future work aims at capturing both, the role of learning and experience for a predator.

The mathematical model that we have developed takes into account the predator prey dynamics with a number of tunable parameters with the predator and its dangerous prey in context. These parameters can be tuned to exact experimental observations. Thus, our study is generalisable to natural predator-prey systems. However, we demonstrate that the eco-evolutionary dynamics of predators and prey are drastically affected when we incorporate innovation dynamics (Fig. 7). An interesting question to ask using an experimental system would be when does selection favour innovative behaviour? While predator-prey interactions themselves can become increasingly complex [49] they are nevertheless a part of community ecology, both biotic and abiotic [50, 51]. In a large, co-evolving food web, analysing whether ecological interactions that are antagonistic or mutualistic promote innovative behaviour would be a study of great importance.

We conclude that incorporating innovation dynamics, together with studies of personality and experience, into traditional evolutionary dynamics will help move current eco-evolutionary theory towards the regimes of an extended evolutionary synthesis applicable across scales of organisation [21].

## Acknowledgements

CSG acknowledges funding from the Max Planck Society. AEW acknowledges funding from Massey University.

## SI.1 Supplementary material

### Simple and complex predators

Herein we develop the dynamics of a safe prey, a dangerous prey and a simple predator (single core tactic) and a complex predator (single novel tactic). How the complex predator arises is not dealt with herein but discussed in the main text.

The density of the safe prey is denoted by *x*_*s*_ and that of the dangerous by *x*_*d*_. The growth rates of the two prey types are *g*_*s*_ and *g*_*d*_, their palatabilities *π*_*S*_ and *π*_*D*_ and the eventual different response curves (*ϕ*_*S*_ and *ϕ*_*D*_) for the safe and dangerous prey respectively. When interacting with a dangerous prey, the predator can get hurt (and eventually die). Dangerous prey is not predators of the predators, so the injury encounter does not increase their net growth rate. The rate at which the predator density can reduce when interacting with the dangerous prey is captured by *ϕ*_*I*_. Taken together the new set of equations is given by,

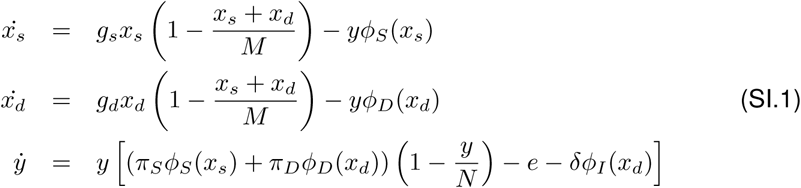

where *M* is the carrying capacity of the prey species and *N* is that of the predators. Now that we have a dangerous prey in the population we turn to the object of this study – a complex predator. A complex predator is the one that can identify when the predator is dangerous and adjust its hunting tactic in such a way that the costs due to injury are minimised and the success rate is better than a näive predator Φ_*D*_ *> ϕ*_*D*_. Thus, now we have the simple predator as *y*, the complex predator *z*,

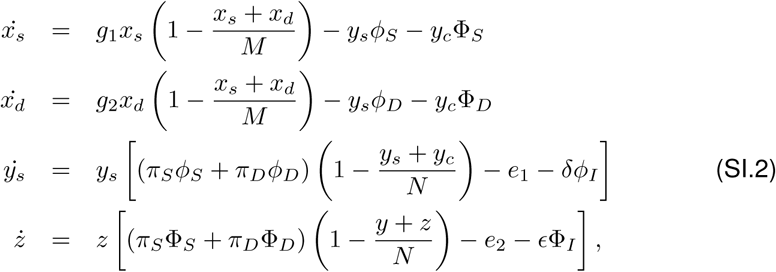

where the response functions are written without the functional form.

---

**Table SI.1:** List of Symbols. Notation used in the manuscript as well as the parameter values used for illustrations.

|  | Description | Value |
| --- | --- | --- |
| $x_s$ | Density of safe prey | |
| $x_d$ | Density of dangerous prey | |
| $g_1$ | Growth rate of safe prey | 3.5 |
| $g_2$ | Growth rate of dangerous prey | 0.1 $g_1$ |
| $M$ | Total carrying capacity of prey | 20 |
| $y_i$ | Density of simple strategy predator $i$ | |
| $z_i$ | Density of complex strategy predator $i$ | |
| $N$ | Total carrying capacity of predator | 20 |
| $\phi_S$ | Core tactic prey capture rate when hunting safe prey | Eqs. (7) |
| $\phi_D$ | Core tactic prey capture rate when hunting dangerous prey | Eqs. (7) |
| $\Phi_S$ | Novel tactic prey capture rate when hunting safe prey | Eqs. (9) |
| $\Phi_D$ | Novel tactic prey capture rate when hunting dangerous prey | Eqs. (9) |
| $\phi_I$ | Core tactic injury rate when hunting dangerous prey | |
| $\Phi_I$ | Novel tactic injury rate when hunting dangerous prey | |
| $f_i$ | Cumulative Holling response for core tactic $i$ | Eq. (11) |
| $F_i$ | Cumulative Holling response for novel tactic $i$ | Eq. (11) |
| $\mu_{i,j}$ | Rate at which the core tactic $j$ switches to $i$ | |
| $r$ | Rate at which novel tactics are formed | $10^{-4}$ |
| $\theta_{i,j}$ | Returns 1 if the core tactics $i$ and $j$ are compatible | for all $i, j = 1$ |
| $D_{i,j}$ | Contribution of the core tactic $i$ when mixed with $j$ | 0.5 |
| $e_1$ | Intrinsic death rate of core tactic predator | 0.75 |
| $e_2$ | Intrinsic death rate of novel tactic predator | 1.5 $e_1$ |

